# Combination of live attenuated and adenovirus-based vaccines completely protects interferon gamma (IFNγ) knockout mice against pneumonic plague

**DOI:** 10.1101/2024.12.06.627261

**Authors:** Emily K. Hendrix, Jian Sha, Paul B. Kilgore, Blake H. Neil, Ashok K. Chopra

## Abstract

Two live attenuated vaccines (LAVs), LMA and LMP, were evaluated alone or in combination with a trivalent adenoviral vector-based vaccine (Ad5-YFV) for their efficacy and immune responses in wild type (WT) and interferon gamma (IFNγ) knockout (KO) mice in a C57BL/6 background. While LMA and LMP are triple deletion mutants of *Yersinia pestis* CO92 strain, Ad5-YFV incorporates three protective plague immunogens. An impressive 80-100% protection was observed in all vaccinated animals against highly lethal intranasal challenge doses of parental *Y. pestis* CO92. All vaccinated mice generated robust humoral and cellular immune responses. The immunized WT mice showed overall better antibody responses in both serum and bronchoalveolar lavage fluid with much higher percentages of polyfunctional T cell populations. On the other hand, vaccinated IFNγ KO mice displayed better B cell activity in germinal centers with higher percentages of activated antigen specific T cells and memory T cells. In addition, depletion of IFNγ and tumor necrosis factor alpha (TNFα) from immunized WT mice prior to and during infection did not reduce protection against pulmonary *Y. pestis* CO92 challenge. These data demonstrated a dispensable nature of IFNγ in mediating protection by the aforementioned vaccines. This is the first detailed immunogenicity study of two plague LAVs administered either alone or in combination with an Ad5-YFV vaccine in a prime-boost immunization strategy in IFNγ KO mice. Further, by combining advantages of live-attenuated and adenovirus-based vaccines, augmentation of generalized immune responses were observed which could be beneficial in providing long-lasting immunity in the host.

## INTRODUCTION

*Yersinia pestis*, the causative agent of bubonic and pneumonic plague, is a Tier-1 select agent responsible for three of seven major pandemics resulting in over 200 million deaths (*1*). Plague remains a conspicuous disease in many regions of the world, and a 2017-2018 epidemic in Madagascar resulted in 2575 cases and 221 deaths, with an 8.6% Case Fatality Rate (*2*). Notably, 77% of these cases were clinically classified as pneumonic plague, incongruent with the majority of bubonic *Y. pestis* infections (*3*). Pneumonic plague exhibits a rapid disease course and is most often fatal if left untreated (*4*). Vaccination is the most effective approach to prevent plague; however, there are currently no Food and Drug Administration (FDA)-approved plague vaccines (*5*). With the existence and/or development of multiple-antibiotic resistant *Y. pestis* strains (*6*), availability of an efficacious vaccine(s) is direly needed. To date, those vaccines that have entered human clinical trials are based on the fraction 1 capsule-like antigen (F1) and the low calcium response V antigen (LcrV) located at the needle tip of the type 3 secretion system (T3SS) (*7–17*). While humoral immune responses are studied in these trials, very few evaluated cellular immunity. Further, there are reports of naturally occurring F1-negative *Y. pestis* strains that remain completely virulent (*18, 19*), making anti-F1 antibodies inconsequential in protection against such *Y. pestis* strains. Likewise, five variants of LcrV have been identified (*20*), offering no cross-protection to immunized mice against subsequent infection with *Y. pseudotuberculosis*, a predecessor of *Y. pestis* (*20*). Thus, F1V vaccines may not confer complete protection against all plague-causing *Y. pestis* strains.

In this regard, live-attenuated vaccines (LAVs) that generate immune responses to thousands of *Y. pestis* antigens would be beneficial to offset lack of protection afforded by antibodies to F1 and LcrV when infection occurs by F1^-^ or strains of yersiniae harboring LcrV variants. The live-attenuated EV76 *Y. pestis* vaccine strain, which is used outside of the United States in Mongolia, China, and the Former States of the Soviet Union, lacks the pigmentation locus (*pgm*) required for iron acquisition from the host, and is protective against both bubonic and pneumonic plague (*21*). However, this vaccine strain behaves as the wildtype (WT) bacterium in individuals with certain metabolic disorders, such as hemochromatosis (*22–24*). In addition, the EV76 vaccine strain generates strong reactogenic side effects and is not suitable for widespread use (*25*), and hence not approved by FDA. Therefore, it is essential to develop novel plague vaccine candidates and employ various immunization strategies that would be protective against F1^-^ *Y. pestis* strains, those which harbor LcrV variants, and are effective in a wide range of human population, especially in regions of endemicity and during an outbreak or bioterror event.

Our laboratory has previously developed two live-attenuated triple deletion mutants of *Y. pestis* strain CO92 (LMA and LMP) and an adenoviral type 5 vector-based vaccine containing 3 prominent plague antigens (Ad5-YFV). The LAV strains are deleted for genes encoding Braun lipoprotein (<u>L</u>pp), acetyltransferase B (<u>M</u>sbB), and attachment-invasion locus (<u>A</u>il), designated as LMA, or plasminogen activating protease (<u>P</u>la), designated as LMP, respectively. Both LMA and LMP mutants were highly avirulent in mice with no clinical symptoms of disease and cleared rapidly (within 12 to 24 h) while retaining immunogenicity (*26–30*). Importantly, the LMA mutant remained avirulent, unlike EV76, during iron overload conditions (*31*). Further, the LMA mutant was well-tolerated in Rag1 KO mice lacking mature B and T cells, showing its promise for use in immunocompromised individuals (*31*). Consequently, these mutants have been excluded from the Centers for Disease Control and Prevention (CDC) select agent list (https://www.selectagents.gov/sat/ exclusions/hhs.html) and recently gained approval from NIH to be used at a lower biosafety level, BSL-2.

Ad5-YFV is a replication-deficient adenovirus type 5 vector-based vaccine containing genes for three plague antigens: *Yersinia* secretion protein F (<u>Y</u>scF), which forms the barrel structure of the T3SS needle, <u>F</u>1, and Lcr<u>V</u> (*32–34*). Addition of YscF circumvents inability of the Ad5-LcrV monovalent vaccine to provide 100% protection against pneumonic challenge in mice with non-capsulated *Y. pestis* CO92 strain (F1^-^ isogenic mutant), as well as against bubonic plague when subjected to challenge with parental *Y. pestis* CO92 (*35*). The Ad5-YFV vaccine was also 100% protective in Cynomolgus macaques (*18*). In addition, heterologous immunization with LMA or LMP and Ad5-YFV vaccines, one dose of each, stimulated robust humoral and cell-mediated immune responses and provided complete protection to mice against lethal challenge with parental *Y. pestis* CO92 and its F1^-^ mutant (*31*). Importantly, increased IFNγ production and overall higher percentages of IFNγ positive T cells were observed in immunized animals irrespective of receiving either LAVs and Ad5-YFV individually or in combination (*31*).

It has been shown that IFNγ plays an integral role in bacterial and viral infections, including *Y. pestis*, by inhibiting pathogen replication while promoting macrophage activation (*36–38*). Several studies have also implicated the importance of IFNγ in F1V subunit plague vaccine mediated protection (*39, 40*). Importantly, inherited IFNγ deficiency due to genetic mutations in IFNγ or its receptors suffer from Mendelian susceptibility to mycobacterial disease (MSMD). This is characterized by a predisposition to severe disease caused by weakly virulent mycobacteria, including the *Mycobacterium bovis* Bacille Calmette-Guérin (BCG) vaccine and environmental mycobacteria(*41, 42*). Therefore, in this study, we further evaluated the safety and immunogenicity of our two LAVs either alone or in combination with Ad5-YFV in both wild type (WT) and IFNγ knockout (KO) mice. Our results showed that our vaccines were well tolerated in IFNγ KO mice without any adverse effects. Although distinct immune responses to vaccination were observed in WT and IFNγ KO mice, all immunized mice were completely protected against lethal pneumonic challenge. Furthermore, depletion of IFNγ and tumor necrosis factor alpha (TNFα) in vaccinated WT mice prior to and during infection did not attenuate protection, indicating a dispensable nature of IFNγ in protection provided by our aforementioned vaccines. This is the first detailed immunogenicity study of two plague LAVs either alone or in combination with Ad5-YFV in IFNγ KO mice.

## RESULTS

### Protection of vaccinated animals against pneumonic plague

WT and IFNγ KO mice were immunized either with LMA or LMP individually (homologous) or in various combinations with Ad5-YFV (heterologous) in a 2-dose prime-boost regimen 21 days apart. The LAVs were administered by the intramuscular (i.m.) route, while Ad5-YFV was instilled by the intranasal (i.n.) route (**Fig. 1A**). No local or systemic adverse effects, such as injection site redness, ruffed fur, and weight loss, were observed in either WT or IFNγ KO mice during vaccination. Roughly 3-weeks post boost vaccination, spleens were excised and bronchoalveolar lavage fluid (BALF) harvested for immunological analysis. On day 53, a separate cohort of mice were i.n. challenged first with 25 LD_50_ of *Y. pestis* CO92 where 1 LD_50_ is 280 colony forming units [CFU] in C57BL/6 mice (*43*). As shown in **Fig. 1B and C**, all vaccinations, regardless of the order or combination of administration, provided 100% protection to both WT and IFNγ mice. As expected, 100% lethality in control mice was observed within 3-4 days post infection (p.i.). Seven days post initial challenge, surviving vaccinated mice along with age-matched control mice were re-challenged i.n. with 10,000 LD_50_ of *Y. pestis* CO92 and observed over a 21-day period. All vaccinations provided 100% protection to WT mice (**Fig. 1D**). However, in both homologous vaccination groups, the protection dropped to 80% in IFNγ KO mice (**Fig. 1E**). All control mice succumbed to infection by day 4. The purpose of rechallenge at an overwhelming dose of *Y. pestis* CO92 was to glean level of protection of immunized mice from recurring infection particularly during an early phase of immune response.

**Fig. 1.**
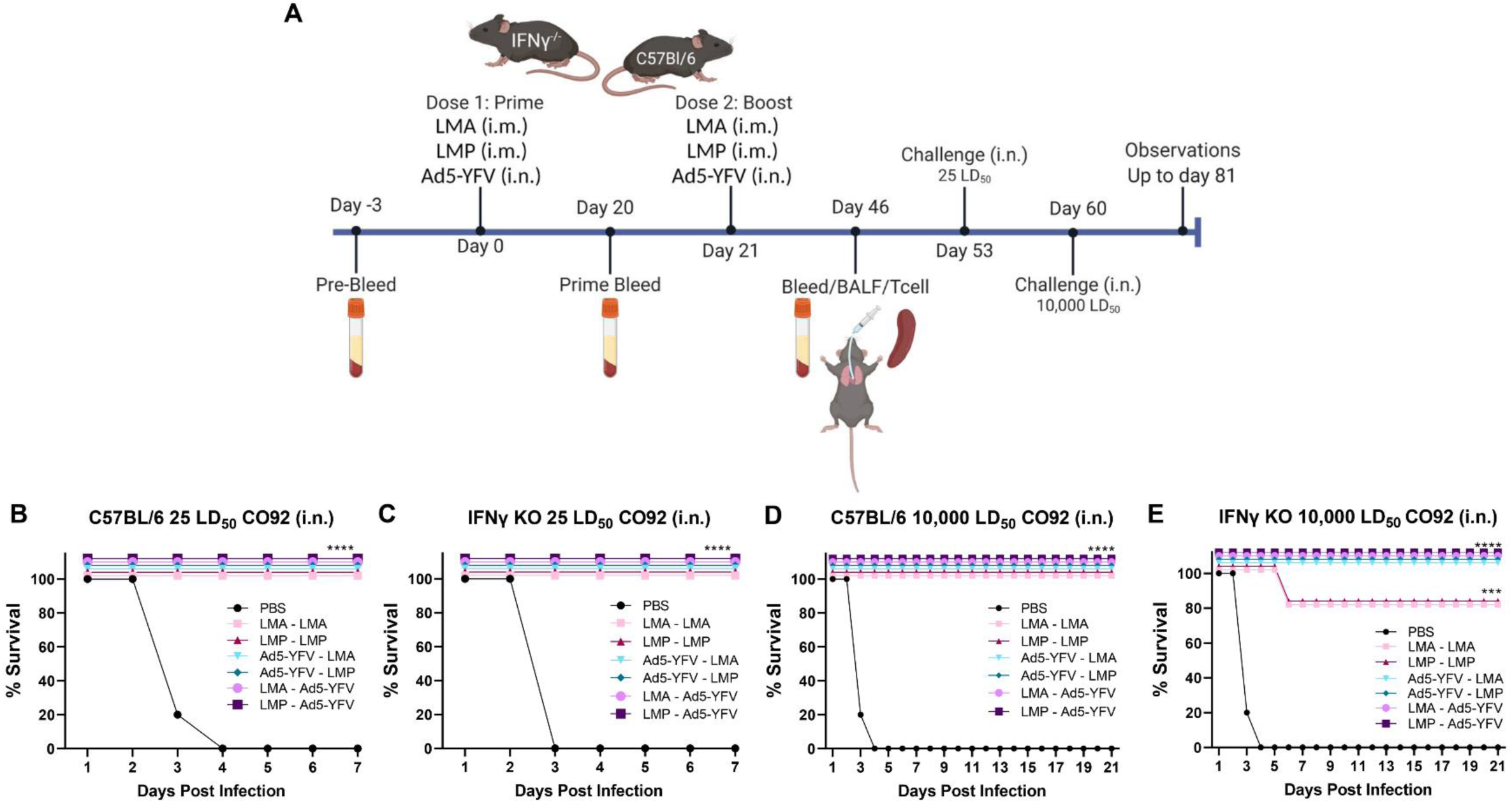
Vaccination protected both WT and IFNγ KO mice against lethal pulmonary *Y. pestis* challenge. Mice (n=10 per group) were immunized with LMA, LMP, or Ad5-YFV in a 2-dose regimen, 21 days apart, in various orders and combinations by either i.n. or i.m. route. Mice receiving PBS were used as controls. The experimental time course is illustrated in panel **A**. Approximately 4 weeks after completion of the vaccination schedule, mice were i.n. challenged with 25 LD_50_ of *Y. pestis* CO92 (**B and C**). Seven days post the initial challenge, surviving vaccinated mice and age-matched naïve control mice were re-challenged i.n. with 10,000 LD_50_ of *Y. pestis* CO92 (**D and E**) and observed for morbidity and mortality for 21 days. Kaplan-Meier analysis with log-rank (Mantel-Cox) test was used to analyze survival. Asterisks represent statistical significance of vaccinated groups compared to naïve control mice. ***, *p* < 0.001; ****, *p* < 0.0001.

### Serum and mucosal antibody responses in vaccinated mice

Serum was collected from immunized and control mice on day 46 prior to *Y. pestis* CO92 challenge (**Fig. 1A**). Both vaccinated WT and IFNγ KO mice had notable increases in IgG titers to recombinant F1V (rF1V) fusion protein over naïve serum (pre-bleed) as well as control animals that received phosphate-buffer saline (PBS) irrespective of the vaccine combination administered (**Fig. 2A**). IgG titers increased between the prime and the boost vaccination by up to 1 log. Compared to WT mice, a relatively lower IgG titer was noticed in IFNγ KO mice with homologous LMP vaccination, while relatively higher IgG titers were exhibited in IFNγ KO mice vaccinated with Ad5-YFV – LMA, Ad5-YFV – LMP, and LMP – Ad5-YFV (**Fig. 2A**). The highest F1V IgG titers were noted in both WT and IFNγ KO mice vaccinated using a prime-pull strategy in which animals received an initial parenteral vaccination of LAVs followed by Ad5-YFV mucosal immunization (lilac and plum bars, **Fig. 2A**). We also examined serum IgG antibody isotypes, IgG1 and IgG2c, to assess the T helper 1 (Th1) versus Th2 bias. Overall, vaccinated groups exhibited significantly higher levels of F1V-specific IgG1 and only moderately higher IgG2c compared to PBS control animals indicating a stronger Th2 response in both immunized WT and IFNγ KO mice (**Fig. 2B and C**). It was further observed that vaccinated IFNγ KO mice exhibited an even lower IgG2c response compared to vaccinated WT mice suggesting that knockout of IFNγ further skewed mice to a Th2 response.

**Fig. 2.**
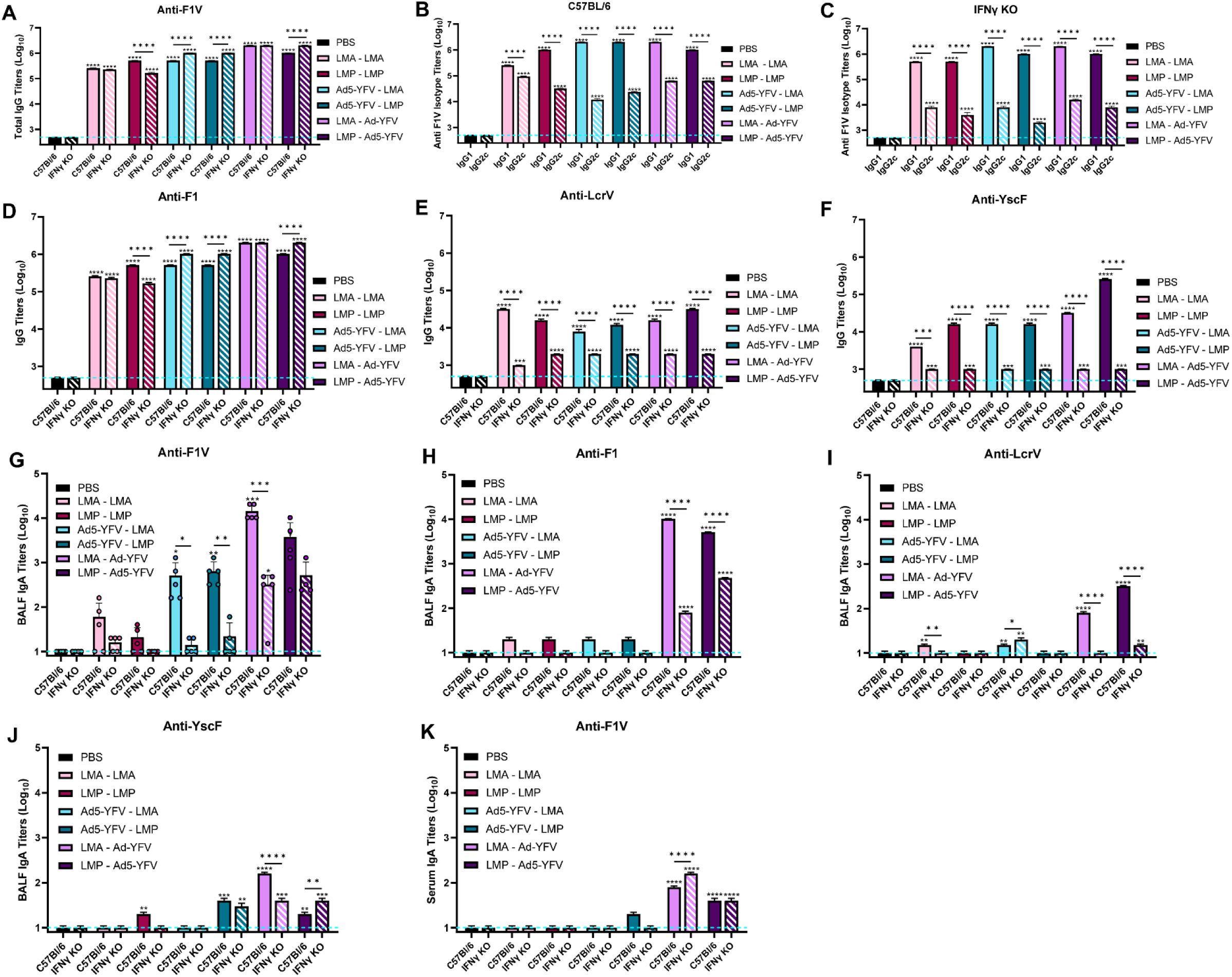
Antibody responses in vaccinated WT and IFNγ KO mice. Animals were immunized, and serum or BALF were collected on day 46 as described in Fig 1A. ELISAs were performed to evaluate the titers: total serum IgG and its isotypes to F1V (**A-C**), or the total serum IgG to individual plague antigens of Ad5-YFV vaccine (**D-F**); BALF IgA titer to F1V (**G**) and individual plague antigens (**H-J**); serum IgA titer to F1V (**K**). Statistical significance was determined by either one-way ANOVA with Tukey’s *post hoc* test (A, D-K) or by two-way ANOVA with Tukey’s *post hoc* test (B and C). The geometric means ± standard deviations are plotted. The blue dashed line represents the baseline values of PBS control samples at the dilution of 1:500. Asterisks with comparison bars represent statistical significance between respective vaccinated groups, while asterisks directly above the bars represent statistical significance of vaccinated groups as compared to naive control mice. *, *p* < 0.05; **, *p* < 0.01; ***, *p* < 0.001; ****, *p* < 0.0001. These data are combined from three independent ELISA assays.

We then further dissected serum IgG antibody titers to 3 individual plague-specific antigens, F1, LcrV, and YscF, which are components of our Ad5-YFV vaccine. Overall, both WT and IFNγ KO mice had significantly higher levels of anti-F1, -LcrV, and -YscF specific IgG than PBS control animals (**Fig. 2D-F**). Increased F1-specific IgG titers over PBS controls reflected the pattern seen for antibody responses to the F1V fusion protein, suggesting the dominant nature of the anti-F1 antibody among the three antibodies measured (**Fig. 2A and D**). Although relatively lower compared to the anti-F1 titer, a significant level of anti-LcrV and anti-YscF specific antibodies were also observed in all vaccinated WT mice compared to unvaccinated control mice, specifically during homologous and prime-pull vaccination strategies (**Fig. 2E and F**). However, IFNγ KO mice mounted significantly lower LcrV- and YscF-specific IgG titers compared to WT mice in response to vaccinations (**Fig. 2E and F**).

To assess mucosal immunity, IgA antibody titers in bronchoalveolar lavage fluid (BALF) were examined. Overall, mice that received heterologous vaccination exhibited significantly higher F1V-specific IgA titers compared to PBS control animals, while immunized IFNγ KO mice mounted a diminished IgA response compared to immunized WT mice (**Fig. 2G**). This trend was more obvious when we examined mucosal IgA antibody titers to the 3 individual plague-specific antigens (F1, LcrV, and YscF). Only mice vaccinated with the prime-pull strategy developed significant IgA antibody responses (**Fig. 2H-J**). A similar phenomenon was observed with serum IgA antibody titers in which the highest F1V titers were found in mice vaccinated with the prime-pull strategy over PBS control mice (**Fig. 2K**).

### Germinal center activity and memory B cells in vaccinated mice

To further assess the humoral immune response, splenocytes were isolated from PBS-administered controls and vaccinated WT and IFNγ KO mice 25 days post boost vaccination (**Fig. 1A**) and stained for germinal center activity (CD3^-^CD138^-^CD19^+^CD38^-^GL7^+^) and memory B cell (CD3^-^CD138^-^CD19^+^GL7^-^CD38^+^IgD^-^) (**Fig. 3A**). Compared to controls, both WT and IFNγ KO mice exhibited increased germinal center B cells after vaccination with either LMP only or the prime-pull strategy; however, vaccinated IFNγ KO animals produced significantly more germinal center B cells than their counterpart WT mice **(Fig. 3B)**. Likewise, WT and IFNγ KO mice also exhibited increased memory B cells only after vaccination with the prime-pull strategy as compared to PBS control animals (**Fig. 3C**).

**Fig. 3.**
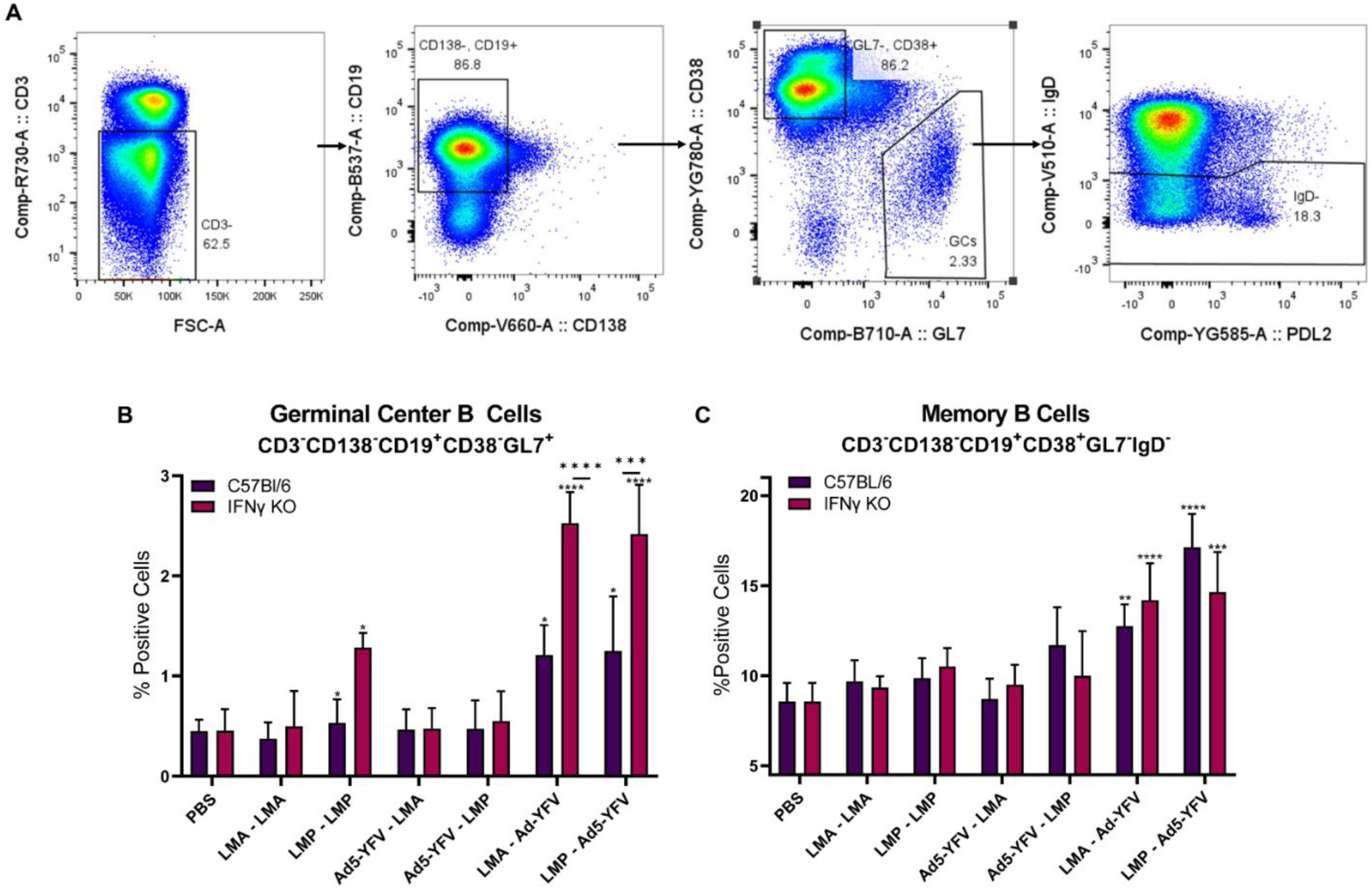
Germinal center B cell activities and memory B cell population in vaccinated WT and IFNγ KO mice. Spleens were collected on day 46 from immunized and naïve control mice (n=5 per group). Splenocytes were surface stained with various B cell markers and subjected to flow cytometry (**A**). The percentage of B cells in germinal center were gated as CD3^-^CD138^-^CD19^+^CD38^-^GL7^+^ (**B**), while memory B cells were identified as CD3^-^CD138^-^CD19^+^CD38^+^GL7^-^IgD^-^ (**C**). The arithmetic means ± standard deviations are plotted. Statistical significance was determined using one-way ANOVA with Tukey’s *post hoc* test. Asterisks with comparison bars represent statistical significance between respective vaccinated groups, while asterisks directly above the bars represent statistical significance of vaccinated groups as compared to naïve control mice. *, *p* < 0.05; **, *p* < 0.01; ***, *p* < 0.001; ****, *p* < 0.0001.

### Vaccination induced cytokine positive T cells

Splenocytes collected post-immunization were stimulated with rF1V and surface stained for anti-CD3, - CD4, and -CD8 followed by intracellular staining for various cytokines (IFNγ, IL-2, TNFα, IL-4, and IL-17). As expected, all vaccinated WT mice had significantly higher populations of IFNγ^+^CD4^+^ and IFNγ^+^CD8^+^cells than control mice, while the stimulation of IFNγ^+^ T cells was abolished in immunized IFNγ KO mice, consistent with the genotype of the KO model (**Fig. 4A and B**). The most significant levels of IFNγ^+^ CD4^+^ and IFNγ^+^ CD8^+^ cells were observed in WT animals vaccinated with LMA-Ad5-YFV. Similarly, all vaccinated WT mice had significantly higher populations of IL-2^+^ or IL-4^+^ CD4^+^ and CD8^+^ T cells than PBS control mice, while these populations were at the basal level in immunized IFNγ KO mice (**Fig. 4C, D, G, and H**). WT mice vaccinated with the homologous LMA elicited the most optimal IL-4 response (**Fig. 4G and H**), and the most significant levels of IL-2^+^ CD4^+^ and CD8^+^ T cells were observed in WT animals vaccinated with LMA-Ad5-YFV, similar to the IFNγ^+^ T cell response (**Fig. 4A and B**).

**Fig. 4.**
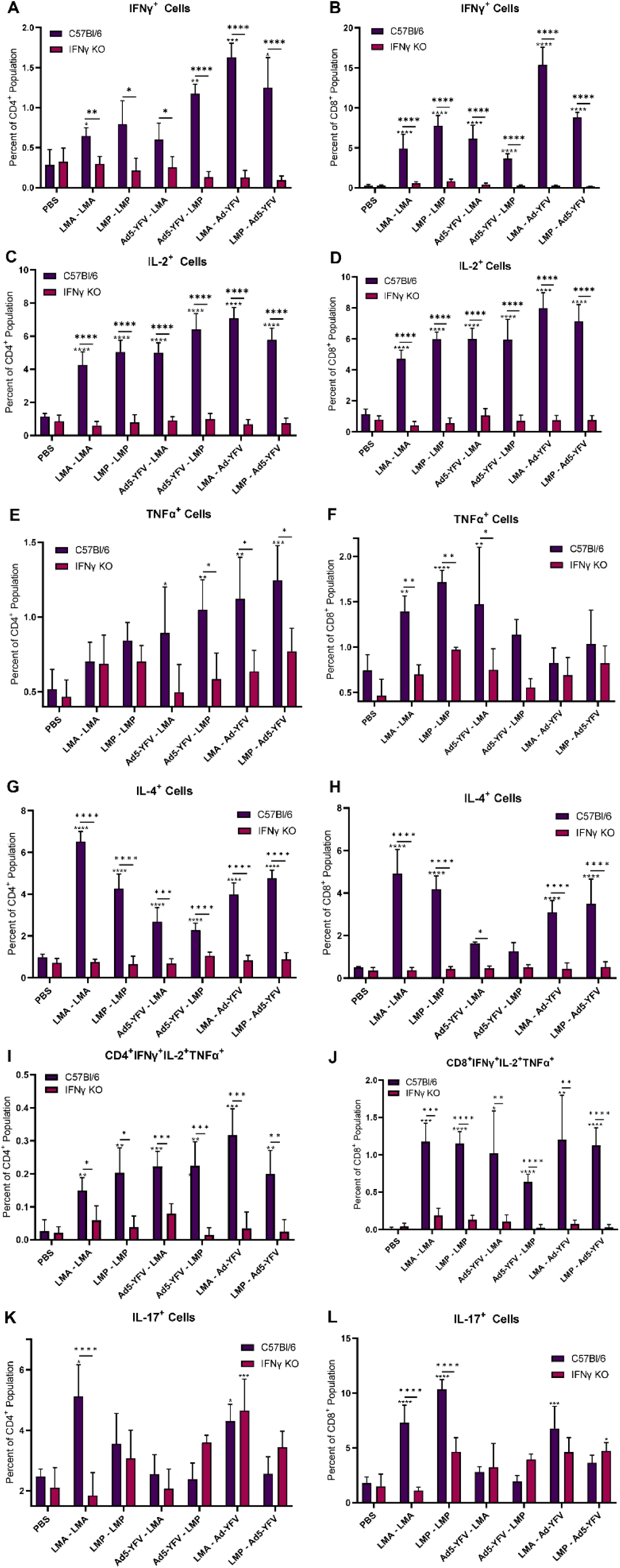
Cytokine positive T cells in vaccinated WT and IFNγ KO mice. Spleens were collected on day 46 from immunized and naïve control mice (n=5 per group). The isolated splenocytes were stimulated with rF1V (100 µg/ml) for 72 h at 37°C followed by additional 4 h with Brefeldin A. Splenocytes were surface and intracellularly stained with various cytokine and T cell markers and subjected to flow cytometry. The percentage of cytokine positive CD4^+^ (**A, C, E, G, I, K**) and CD8^+^ (**B, D, F, H, J, L**) populations were acquired. The arithmetic means ± standard deviations are plotted. Statistical significance was determined using one-way ANOVA with Tukey’s *post hoc* test. Asterisks with comparison bars represent statistical significance between respective vaccinated groups, while asterisks directly above the bars represent statistical significance of vaccinated groups as compared to PBS control mice. *, *p* < 0.05; **, *p* < 0.01; ***, *p* < 0.001; ****, *p* < 0.0001.

TNFα is also a Th1 cytokine and can synergize with IFNγ to mitigate infection(*44, 45*). Compared to PBS control, elevated levels of TNFα^+^ T cells were observed in all immunized WT mice. However, levels were much lower than IFNγ, IL-2 and IL-4, and only reached significant levels for CD4^+^ T cells in heterologous vaccinated groups and of CD8^+^ cells in homologous immunized groups. A basal level of TNFα^+^ T cells was noticed in all vaccinated IFNγ KO mice (**Fig. 4E and F**). Most importantly, polyfunctional T cells positive for all 3 Th1 cytokines (IFNγ, IL-2, and TNF-α) were significantly higher in all immunized WT mice compared to vaccinated IFNγ KO and PBS control mice (**Fig. 4I and J**). Collectively, these results indicated the ability of our vaccines to elicit T cell responses in WT mice, and IFNγ was implicated in the production of IL-2^+^, TNFα^+^, and IL-4^+^ T cells in response to vaccination.

Since IL-17 contributes to cell-mediated defense against pulmonary *Y. pestis* infection (*46*), we examined IL-17^+^ T cell response. Unlike the above Th1 and Th2 cytokine positive T cells, which were at basal or reduced levels in all vaccinated IFNγ KO mice, comparable levels of IL-17^+^ T cells were observed in both WT and IFNγ KO mice (**Fig. 4K and L**). More specifically, WT mice that received homologous LAVs or prime-pull LMA-Ad5-YFV vaccination showed significant increases in IL-17^+^ CD4^+^ and/or CD8^+^ T cells compared to PBS control. On the other hand, IFNγ KO mice immunized with the prime-pull strategy had higher IL-17 responses in either CD4 or CD8 T cells (**Fig. 4K and L**). Interestingly, in homologous LAV vaccination groups, a significantly lower IL-17 response was observed in IFNγ KO mice compared to their WT counterpart, indicating IFNγ might be required for IL-17 production during LAV vaccination (**Fig. 4K and L**).

### Activated antigen-specific T cells

To fully capture the repertoire of antigen-specific T cells elicited by vaccination, we employed a more inclusive, cytokine agnostic approach of an activation-induced markers (AIM) assay. Antigen experienced T cells (CD44^+^) were further gated for CD4 and CD8 activation markers CD134^+^CD25^+^ and CD69^+^CD25^+^, respectively. As shown in **Fig. 5**, generally all vaccinated mice had relatively more antigen-activated T cells than that of unvaccinated control mice. Surprisingly, the vaccinated IFNγ KO mice possessed significantly higher antigen-activated T cells than their corresponding WT counterparts. This contrasted with the Th1 and Th2 cytokine positive T cells (**Figs. 4 and 5**). This phenomenon indicated knockout of IFNγ altered the repertoire of antigen-specific T cells in response to vaccination.

**Fig. 5.**
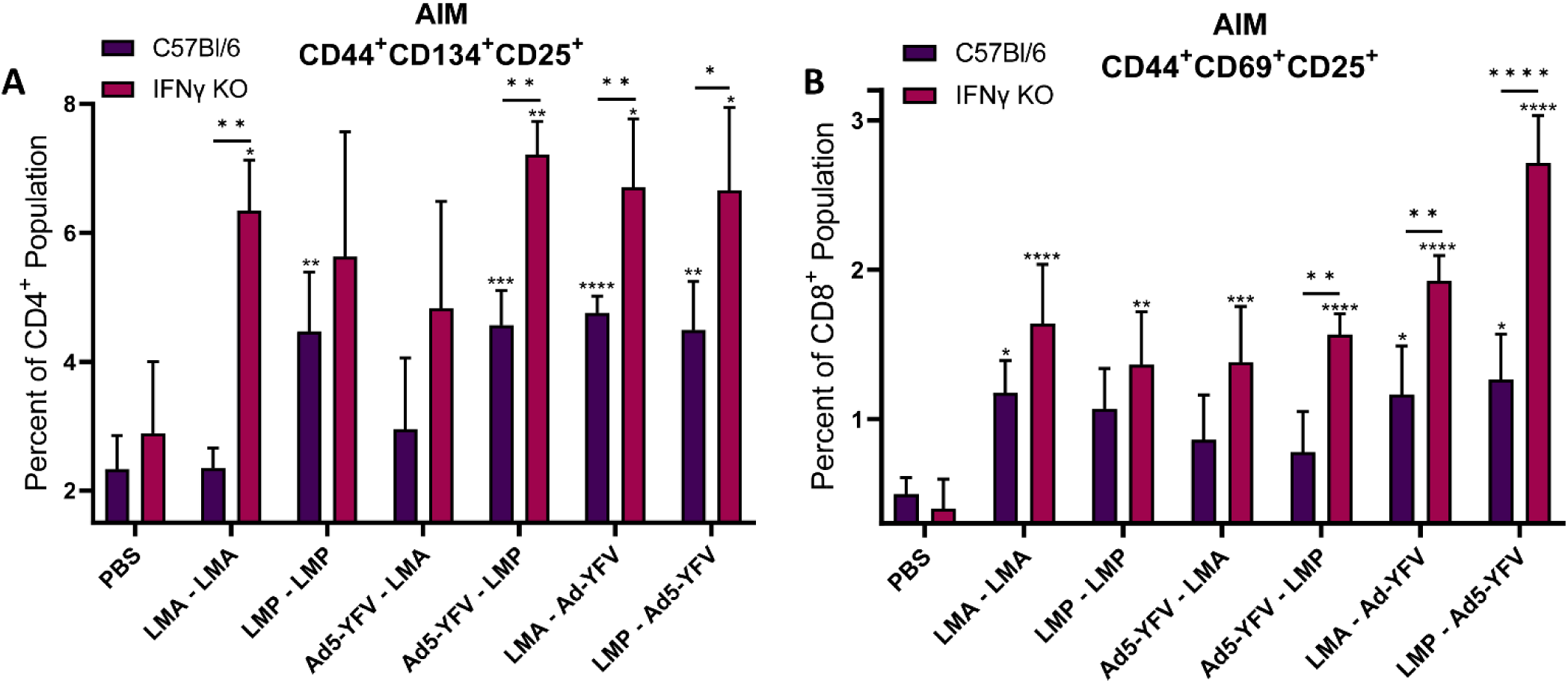
Evaluation of T cell response by using activation induced marker (AIM) assay. Spleens were collected from vaccinated and control mice (n=5 per group) and stimulated with rF1V (100 µg/ml) for 72 h at 37°C followed by additional 4 h with Brefeldin A, as stated in the **Fig. 4** legend. T cells were then harvested and stained with specific T cell AIM markers for flow cytometry analysis. The percentage of positive AIM populations in CD4^+^CD44^+^CD134^+^CD25^+^ (**A**) and CD8^+^CD44^+^CD69^+^CD25^+^(**B**) were acquired. The arithmetic means ± standard deviations are plotted. Statistical significance was determined using one-way ANOVA with Tukey’s *post hoc* test. Asterisks with comparison bars represent statistical significance between respective vaccinated groups, while asterisks directly above the bars represent statistical significance of vaccinated groups as compared to naïve control mice. *, *p* < 0.05; **, *p* < 0.01; ***, *p* < 0.001; ****, *p* < 0.0001.

### Memory T cells in vaccinated mice

Given the importance of memory immunity in protection against pathogen reinfection, we evaluated central (T_CM_) and effector (T_EM_) memory T cell responses. Both T_CM_ and T_EM_ subsets circulate in the blood. T_CM_ cells interact with antigen-specific dendritic cells in primary lymphoid tissues and subsequently expand to acquire effector function. T_EM_ cells migrate into secondary lymphoid tissues and directly interact with pathogens, providing cytotoxic functions(*47*). Compared to PBS control mice, knockout of IFNγ overall significantly increased vaccine-induced central (**Fig. 6A and B**) and effector (**Fig. 6C and D**) memory T cells. Only WT mice vaccinated with homologous LAVs exhibited higher CD8^+^ Central Memory T cells, though still significantly lower than their IFNγ KO counterpart (**Fig. 6B**). These data indicated that deletion of IFNγ altered the profile of memory T cells in response to vaccination.

**Fig. 6.**
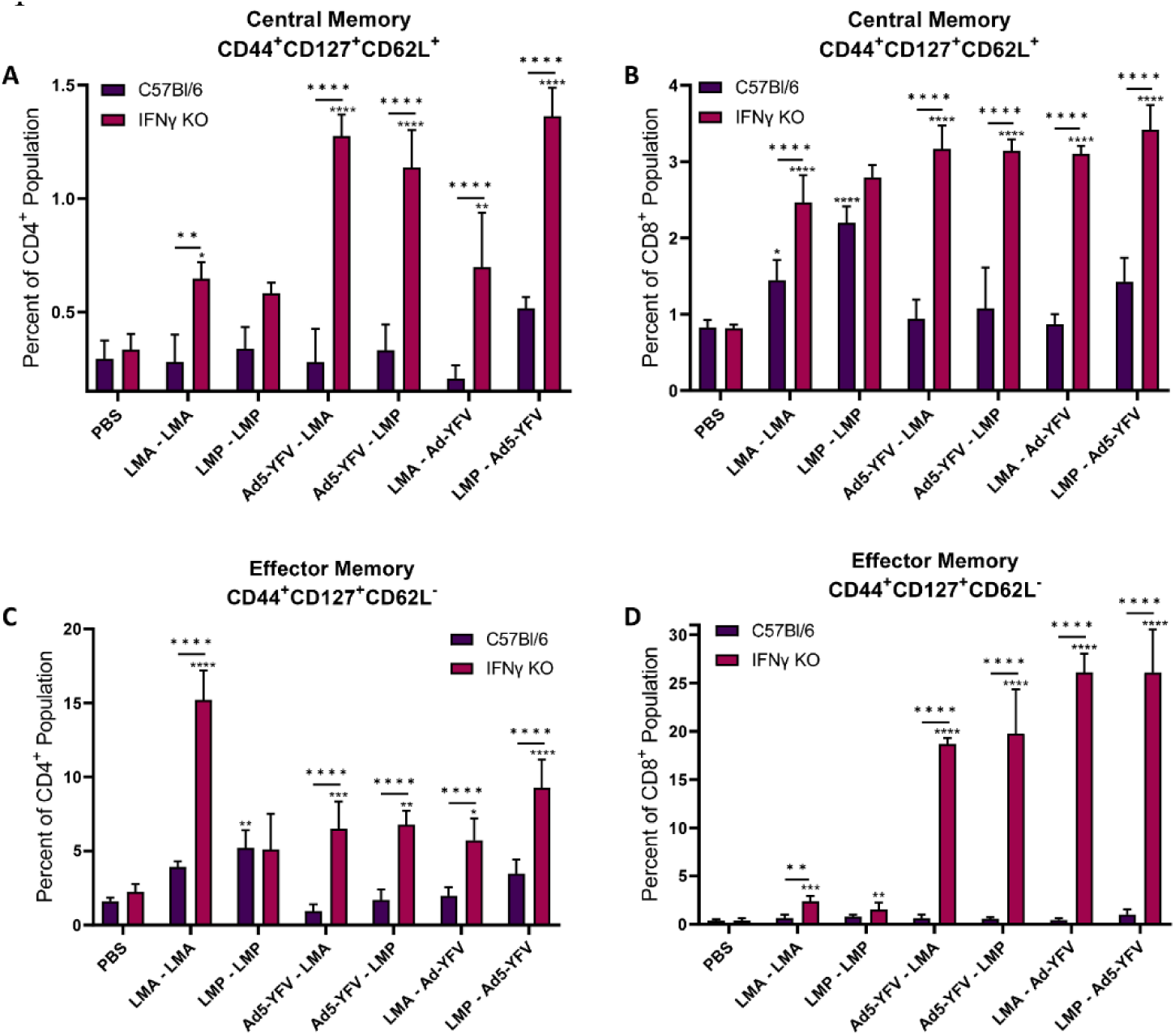
Memory T cell populations in vaccinated WT and IFNγ KO mice. Spleens were collected from immunized and control mice (n=5 per group) and stimulated with rF1V (100 µg/ml) for 72 h at 37°C followed by additional 4 h with Brefeldin A, as stated in **Fig. 4** legend. Cells were then harvested and stained with various T cell markers and subjected to flow cytometry. Central memory T cells (T_CM_) were gated as CD44^+^CD127^+^CD62L^+^ (**A and B**) while effector memory T cells (T_EM_) were identified as CD44^+^CD127^+^CD62L^-^ (**C and D**). The arithmetic means ± standard deviations are plotted. Statistical significance was determined using one-way ANOVA with Tukey’s *post hoc* test. Asterisks with comparison bars represent statistical significance between respective vaccinated groups, while asterisks directly above the bars represent statistical significance of vaccinated groups as compared to naïve control mice. *, *p* < 0.05; **, *p* < 0.01; ***, *p* < 0.001; ****, *p* < 0.0001.

### Protection of vaccinated WT mice upon IFNγ or TNFα depletion

To fully gauge importance of Th1 cytokines, specifically IFNγ and TNFα in vaccine-mediated protection, as well as any off-target effects associated with IFNγ gene knockout, we depleted IFNγ or TNFα in WT mice post vaccination. The prime-pull strategy with 1- or 2-dose regimes were used in these studies as they generated superior immune responses in both our current and previous studies against *Y. pestis* CO92 and its F1^-^ mutant (*35*). In a 2-dose series, LMA or LMP was given i.m. followed by Ad5-YFV i.n. 21 days apart, while in 1-dose regime, a single dose of the above vaccines was provided simultaneously. A simultaneous vaccination strategy could be beneficial in an emergent biothreat or outbreak scenario (*31*). Three weeks post boost vaccination, mice were treated intraperitoneally (i.p.) with mAbs either for IFNγ, TNFα, or IgG control (Bio X Cell, Lebanon, NH) in 3 doses: one day prior, at the time of, and one day post infection. Mice were challenged with 100 LD_50_ of *Y. pestis* CO92 by i.n route. All vaccinated mice survived lethal pneumonic challenge irrespective of mAb treatment received, while 100% of unvaccinated mice succumbed to infection (**Fig. 7A-C**). In addition, all *Y. pestis* bacilli were cleared from lung and spleen tissues of vaccinated mice by day 21 p.i. These data indicated that neither the gene deletion of IFNγ prior to vaccination nor its depletion post vaccination affected the protective efficacy of our vaccines.

**Fig. 7.**
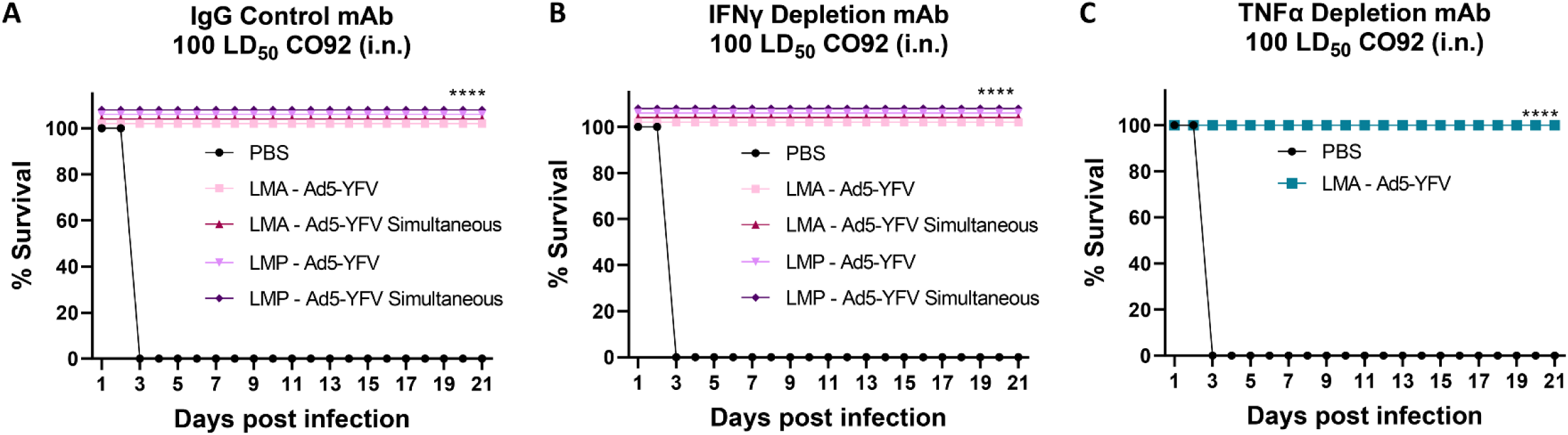
Evaluating the protection of vaccinated WT mice against lethal pulmonary *Y. pestis* challenge upon IFNγ or TNFα depletion. Mice (n=7 per group) were immunized with LAVs (LMA or LMP, i.m.) plus Ad5-YFV (i.n.) in either a 1- or 2-dose regimen. In the 1-dose regimen, both LAV and Ad5-YFV were administered simultaneously. In the 2-dose regimen, LAV and Ad5-YFV were administered 21 days apart. Mice receiving PBS were used as challenge control. Three weeks after completion of the vaccination schedule, the immunized mice were treated with either an IgG control mAb (**A**), depletion mAb for IFNγ (**B**), or depletion mAb for TNFα (**C**) at 1 day before, at the time, and 1 day after 100 LD_50_ of intranasal *Y. pestis* CO92 challenge. Animals’ morbidity and mortality were observed for 21 days. Kaplan-Meier analysis with log-rank (Mantel-Cox) test was used to analyze survival. Asterisks represent statistical significance of vaccinated groups compared to naïve control mice. ****, *p* < 0.0001.

## DISCUSSION

As a central effector of cell-mediated immunity, IFNγ plays a critical role in the recognition and elimination of invading pathogens by amplifying interactions between antigen presenting cells (APCs) and T cells (*36–38*). IFNγ released by Th cells also contributes to activation of effector cells and stimulation of humoral responses (*48, 49*). Abrogation of IFNγ or its receptor (IFNγR) increases susceptibility to certain bacterial infections, such as *Mycobacterium tuberculosis*, *Listeria monocytogenes*, and *Y. pestis*, among others (*50–53*). In fact, the correlation of IFNγ and TNFα with serotherapy- and vaccination-mediated protection against pneumonic plague has been demonstrated and suggested to be used as surrogate assays for plague vaccine efficacy in humans (*40, 53*).

The current study is aimed to elucidate the impact of IFNγ on the efficacy of our live-attenuated and adenovirus-based plague vaccines under either homologous or heterologous vaccination strategies. Our data showed full protection against pulmonary *Y. pestis* challenge in all vaccinated WT and IFNγ KO mice with strong humoral and cellular immune responses; however, each mouse type and vaccination strategy displayed their own distinctive immune characteristics.

In murine models, C57BL/6 mice are considered the prototypical Th1-type strain. Surprisingly, both our vaccinated WT and IFNγ KO mice exhibited significantly higher levels of IgG1 over IgG2c, and IFNγ KO mice exhibited an even lower IgG2c compared to WT mice suggesting that knockout of IFNγ further skews mice to a Th2 response. This Th2 bias was also partially reflected in the cytokine profile of CD4^+^ T cells, especially in homologous LAV vaccinated WT groups. For example, the percentage of Th2^+^ (IL-4) CD4^+^ T cells was generally higher than Th1^+^ (IFNγ or TNFα) CD4^+^ T cells. However, more balanced Th1^+^ and Th2^+^ CD4^+^ cells were observed in heterologous, specifically prime-pull, immunized groups of mice (**Fig. 4**), possibly due to the adjuvant nature of the adenovirus vector to promote a Th1 response (*32*). In our previous studies, we observed a Th1 or balanced immune response in outbred Swiss Webster mice receiving either homologous (LAVs or Ad5-YFV) or heterologous (LAVs in combination with Ad5-YFV) vaccination (*31, 35*). This difference may be related to different mouse strains used. Studies have shown that mucosal immunization with LAVs provide better protection against *Francisella tularensis* (*Ft*) in outbred Swiss Webster mice compared to inbred C57BL/6 due to enhanced T cell immunity (*54*). It has also been demonstrated that C57BL/6 mice are less protected than BALB/c mice against virulent type A *Ft* challenge, attributed to waning T cell immunity in C57BL/6 mice (*55*). Furthermore, C57BL/6 mice favor the development of a Th2 phenotype rather than the more protective Th1 response in the lungs in response to BCG vaccination or *Cryptococcus neoformans* infection(*56, 57*). Nevertheless, our vaccines provided complete protection irrespective of the type of Th response mounted in either Swiss Webster or C57BL/6 mice.

IgA, the predominant antibody isotype in mucosal tissue, is an integral component of pathogen defense at mucosal sites (*58, 59*). Secretory IgA, the most prevalent form of IgA in mucosal secretions, is more efficient than IgG in the serum for protection against respiratory pathogens (*60, 61*). In our study, heterologous vaccination of mice, especially by the prime-pull strategy, significantly increased BALF IgA titers compared to control animals, while homologous LAV vaccination was less successful (**Fig. 2**). Interestingly, BALF IgA levels in IFNγ KO mice were significantly diminished compared to WT animals, indicating that IFNγ is needed for local mucosal plasma cells to produce IgA. Studies have shown abrogation of IgA leads to lower levels of pulmonary IFNγ, regardless of antigen-specific IgM and IgG levels (*58, 62*). However, it has also been found that fecal IgA anti-LPS antibody was almost fourfold lower in WT than in IFNγ KO mice in response to *Salmonella* Typhimurium infection (*52*). Therefore, the impact of IFNγ on secretory IgA production may depend on the type of mucosal tissue and pathogens used. Interestingly, the elicited serum IgA in our study was only observed in mice that received prime-pull vaccination and their levels were comparable in both WT and IFNγ KO mice (**Fig. 2K**). It has been reported in a study utilizing IgA and IgM deficient mice (polymeric IgR-deficient, pIgR^-/-^) that secretory IgA could be dispensable in protecting mice against pneumonic plague (*63*). However, pIgR^-/-^ mice exhibit immunological differences in addition to IgA compared to WT mice (*64*). Furthermore, the role of secretory IgA against pneumonic plague could be masked by the high level of protective serum IgG (*65*). Therefore, the role of secretory IgA during pneumonic plague is still not fully elucidated and requires further investigation.

Generation of protective antibodies relies on production of a robust response within germinal centers (GCs) of lymphoid organs (*66*). GCs are transient structures that form within peripheral lymphoid organs in response to T cell-dependent antigens, where antigen-activated B cells undergo somatic hypermutation and selection driving antibody affinity maturation and long-lived memory B cell and plasma cell differentiation (*66–70*). In our study, significantly higher GC-derived B cells were only observed in either LMP homologous or prime-pull heterologous vaccination approaches when compared to unvaccinated animals, which were further augmented in IFNγ KO mice under these vaccination strategies (**Fig. 3B**). Similarly, we noticed an increase in memory B cells only in the same prime-pull vaccination groups, although these were more uniform in response between WT and IFNγ KO mice (**Fig. 3C**). Our data indicated that GC activities were not only influenced by the nature of vaccines but also by the vaccination strategies, and IFNγ may play a role in dampening GC response to plague antigens.

Studies have shown that GC activity and B cell maturation are associated with protective antibody responses against *Plasmodium* preerythrocytic infection elicited by vaccination with *Plasmodium yoelii* circumsporozoite protein (*71*). Similarly, enhanced GC T follicular helper (Tfh) and B cell responses were correlated with long-lasting and high F1V-specific antibody titers and more robust antibody recall responses to F1V re-exposure in mice vaccinated with a plague subunit F1V vaccine in combination with CpG ODN bound to alhydrogel (*66*). Interestingly, in our study, there is no difference in serum antibody titers to all three plague antigens (F1, LcrV and YscF) within the same type of immunized mice (either WT or IFNγ KO) irrespective of the vaccination strategy. Furthermore, antibody titers to LcrV and YscF were significantly lower in immunized IFNγ KO mice compared to immunized WT mice, although antibody titers to the F1 antigen were generally comparable in these mice. The decorrelation between antibody titers and GC activities could be explained by our measurement of the overall response to a variety of antigens from LAVs in GCs, while the antibody titers specifically measured F1, LcrV and YscF. In addition, GC activities corelated better with neutralization antibodies and memory responses (*66, 71*). Therefore, further study of GC activities for specific antigens, measuring neutralization antibody titers, and long-term protection may help us fully understand the role of GC activities in our vaccinated animals.

As *Y. pestis* is a facultative intracellular bacterium, cell-mediated immunity is integral to defense against this pathogen. In response to stimulation, T lymphocytes undergo complicated patterns of differentiation from uncommitted precursors to highly competent effector cells of at least two distinct subsets, Th1 and Th2, which are characterized by both their function and cytokine profiles (*72*). IFNγ and IL-4 are prototypic Th1 and Th2 cytokines, respectively. The cytokine milieu during and after the process of antigen recognition has been shown to be a critical determinant for T cell differentiation (*73*). It is well documented that IL-12 and IL-4 are major determinants of the Th commitment process (*74, 75*). However, the roles that IFNγ plays remain controversial (*72*).

For example, IFNγ was found to be required for stabilization of the Th1 phenotype in several rounds of *in vitro* priming of CD4^+^ T cells (*76*). Similarly, inbred C57BL/6 mice resistance to *Leishmania major* infection is Th1-dependent, and absence of IFNγ results in skewing to a Th2 response and failure to successfully control the infection(*77–79*). On the other hand, in the absence of IFNγ receptors, Th1 responses still develop in genetically *L. major* resistant 129/Sv/Ev mice with no evidence for expansion of Th2 cells(*80*). In fact, it has been shown that IFNγ and IL-4 synergize *in vitro* to enhance CD4 cells to produce IL-4 and their Th2 differentiation. Furthermore, *in vivo* priming for IL-4 production in IFNγ-deficient mice revealed that complete absence of IFNγ during priming leads to less optimal Th2 differentiation, while addition of IFNγ during priming enhances Th2 polarization to levels significantly higher than in wild-type animals (*72*). A similar trend was observed in our study as the percentage of IL-2^+^, TNFα^+^, or IL-4^+^ CD4^+^ and CD8^+^ T cells were significantly lower in immunized IFNγ KO mice compared to immunized WT mice, suggesting abrogation of IFNγ reduces both Th1 and Th2 responses (**Fig. 4**).

In addition to Th1 and Th2, Th17 cells play an integral role in protection against extracellular pathogens, especially in mucosal tissues, by stimulating chemokine release from macrophages, endothelial, and epithelial cells (*81*). In fact, IL-17 production is an important correlate of protection against plague in the absence of protective antibodies (*31, 46*). Both Th1 and Th17 responses are well known to mediate autoimmune inflammatory diseases. It has become clear that the relationship between Th1 and Th17 lineages is much more intertwined and complex than was initially appreciated. While there is clear evidence of counter regulation, there is also increasing evidence of cooperation, and even dependency, in affecting pathology(*82*). Unlike the influence of IFNγ on Th1 and Th2 responses, in our study, the Th17 response was generally comparable between corresponding WT and IFNγ KO groups, except for groups immunized with homologous LAVs, in which significantly higher levels of IL-17 were observed for CD4^+^ and/or CD8^+^ T cells in WT mice. Compared to unimmunized mice, Th17 responses were elicited only in mice receiving homologous LAVs or heterologous vaccines with LAVs as the first dose (**Fig. 4**). These findings agree with our previous study for LMA vaccination that elicited a significant induction of IL-17^+^ CD4^+^ T cells in outbred mice (*31*). Therefore, our data indicated that Th17 response was not only influenced by IFNγ, but also by the nature of our vaccines, which reflects the complexity of the relationship between Th1 and Th17.

The diminished production of both Th1 and Th2 cytokine positive T cells in IFNγ KO mice was unexpected as all vaccinated IFNγ KO mice were completely protected from lethal *Y. pestis* challenge. As such, we further evaluated the T cell response by using an activation induced marker (AIM) assay, which is based on upregulation of T cell receptor-stimulated surface markers (*83*). Importantly, AIM can identify activated T cells even if they do not express sufficient cytokines to be measured by traditional intracellular staining (ICS) methods (*84*). Indeed, significantly higher activated antigen specific T cells were observed in all vaccinated IFNγ KO mice than in immunized WT mice (**Fig. 5**). Furthermore, immunized IFNγ KO mice also exhibited increased memory T cell levels over both control and WT mice (**Fig. 6**), indicating a strong cellular immune response was mounted. Compared to homologous vaccination, enhanced memory T cell responses were associated with heterologous immunization. These findings agree with previous studies that anti-IFNγ antibodies could improve immunogenicity of recombinant adenovirus vectors by promoting development of memory CD8 T cells (*85*), and lack of IFNγ-receptor signaling in CD8^+^ T cells promotes long-lived memory T cell formation in response to weak, but not strong, TCR agonists (*86*).

In summary, the immunized WT mice showed overall better antibody responses to LcrV and YscF in both serum and BALF with much higher percentages of cytokine positive T cells, including polyfunctional T cells. On the other hand, vaccinated IFNγ KO mice displayed better B cell activity in GCs with higher percentages of activated antigen-specific T cells and memory T cells. In addition, depletion of IFNγ and TNFα from immunized WT mice did not reduce protection against pulmonary *Y. pestis*challenge. These data clearly suggest a dispensable nature of IFNγ in vaccine-mediated protection with our vaccines, which is possibly due to the optimal antibody response elicited in immunized mice that overrides the need for IFNγ and TNFα (*40*).

Among the immunization strategies used in this study, the prime-pull strategy showed optimal IgA production and GC B cell activities, a balanced Th1/Th2 response, and increased level of activated antigen specific T cells along with superior memory B and T cell populations. Therefore, both our vaccines and associated vaccination strategies are very promising and should be further evaluated in larger animal models, especially in Cynomolgus macaques and African green monkeys. Our future studies will also focus on the prime-pull strategy with our vaccines to study long term protection and efficacy against non-capsulated *Y. pestis* CO92.

## MATERIALS AND METHODS

### Bacterial strains and vaccines

The fully virulent human pneumonic plague isolate, *Y. pestis* CO92, was obtained from BEI Resources (Manassas, VA). The live-attenuated vaccine candidates LMA and LMP are triple deletion mutants (Δ*lpp*Δ*<u>m</u>sbB*Δ*<u>a</u>il* [LMA] and Δ*lpp*Δ*<u>m</u>sbB*Δ*<u>p</u>la* [LMP]) of CO92 deleted for genes encoding Braun lipoprotein (Lpp), acetyltransferase B (MsbB), and attachment-invasion locus (Ail) or plasminogen activating protease (Pla), respectively(*26, 30*). The replication-deficient human adenoviral type 5 vector-based vaccine Ad5-YFV includes genes for three plague antigens: YscF (T3SS barrel), F1 (capsule), and LcrV (T3SS needle) (*32*). All studies utilizing *Y. pestis* were performed in Tier 1 select agent BSL3 laboratories at UTMB in the Galveston National Laboratory (GNL)-Keiller complex in Galveston, Texas. The Ad5-YFV vaccine was purified at the Baylor College of Medicine Vector Development Laboratory and by our company partner in collaboration with Lonza, Houston, TX, under good laboratory practice conditions. The vaccine preparations were aliquoted at 1×10^12^ v.p./mL and stored at -80°C. These preparations were used for all studies in mice using ABSL3 facilities of GNL-Keiller complex (*32, 35*). The LMA and LMP vaccines were prepared in the GNL biosafety level 3 (BSL3) and aliquoted (500µL, 1̴×10^9^CFU/ml) with 25% glycerol and stored at -80°C. Titers of the vaccines were routinely confirmed before and after each inoculation. These preparations were used for all studies in mice (*26, 30*).

### Animals

Female inbred C57BL/6 wild-type (WT) and IFNγ knockout (KO; B6.129S7-*Ifng^tm1Ts^*/J) mice (6 weeks) were purchased from the Jackson Laboratory (Bar Harbor, ME). All studies were ethically performed under an approved Institutional Animal Care and Use Committee (IACUC) protocol.

### Vaccination and challenge studies

Either C57BL/6 (WT) or C57BL/6 IFNγ KO mice were immunized with two doses of LMA or LMP intramuscularly (i.m.), or combinations of Ad5-YFV intranasally (i.n.) and LMA or LMP i.m. All vaccines were administered 21 days apart. Mice receiving PBS were used as controls. Vaccination dose for Ad5-YFV was 5.5×10^10^ PFU/50µL and for LMA or LMP was 2×10^6^ CFU/50µL.

Mice (n=10 per group) were retro-orbitally bled and serum samples pooled prior to vaccination and around 21 days after each immunization. On day 46, spleens and BALF were collected individually from 5 mice in each group for immunological analysis. The remaining 5 mice from each group were then challenged with 25 LD_50_ of *Y. pestis* CO92 on day 53. Seven days post initial challenge, surviving mice and naïve age-matched control mice were re-challenged with 10,000 LD_50_ of *Y. pestis* CO92.

### Analysis of antibody titers

Antibody titers were measured via indirect enzyme-linked immunosorbent assays (ELISAs). Briefly, MaxiSorp Nunc ELISA plates (Fisher Scientific, Hampton, NH) were coated with 100 ng of either recombinant fusion protein (rF1V) (BEI Resources) or individual antigens rF1, rLcrV, or rYscF in carbonate buffer (total 100 µL) overnight at 4°C. Free antigens were removed by three washes with Dulbecco’s PBS (DPBS) containing 0.05% Tween 20. Plates were blocked with 1% bovine serum albumin (BSA) (Sigma Aldrich, St. Louis, MO) in DPBS for 2 h at room temperature. Serum and BALF were initially diluted with DPBS and then 2-fold serially diluted and incubated for 1 h at room temperature. Plates were washed 3 times with DPBS+0.05% Tween 20 and then horseradish peroxidase (HRP)-conjugated secondary anti-mouse antibodies for IgG, IgG1, IgG2c, or IgA (Southern Biotech, Birmingham, AL) diluted to 1:8000 were added for 1 h at room temperature. Plates were washed three times and 100 µL 3,3’,5,5’-tetramethyl-benzidine (TMB) substrate was added for up to 10 min at room temperature. Development of the colorimetric reaction was stopped using 2N H_2_SO_4_ and absorbance measure at 450nm using a VersaMax tunable microplate reader (Molecular Devices, San Jose, CA). Total IgG, IgG1, IgG2c, and IgA were measured in triplicate.

### Flow cytometry analysis. T cell phenotypes

Spleens that were collected from control and immunized mice, and single-cell suspension was made in RPMI 1640 medium. Splenocytes were seeded into 96-well tissue culture plates at a density of 2.0×10^6^ cells per well and stimulated with 100 µg/mL of rF1V antigen (BEI Resources) for 3 days at 37°C in a 5% CO_2_ incubator. After 3 days of stimulation, 1X Brefeldin A was added for additional 4 h. Splenocytes were then harvested and blocked with anti-mouse CD16/32 antibodies (Biolegend, San Diego, CA) followed by staining with fixable viability dye eFluor 780 (Invitrogen, Waltham, MA), Alexa Fluor 700 anti-mouse CD3 (Biolegend), Brilliant Violet 785 anti-mouse CD4 (Biolegend), FITC anti-mouse CD8 (Biolegend), Brilliant Violet 510 anti-mouse CD44 (Biolegend), Brilliant Violet 711 anti-mouse CD62-L (Biolegend), PE anti-mouse CD127 (Biolegend), Brilliant Violet 605 anti-mouse CD25 (Biolegend), and APC anti-mouse CD134 (Biolegend). Cells were then permeabilized for intracellular staining with the Foxp3/Transcription Factor Staining Buffer Set (eBioscience, San Diego, CA) at 4°C overnight and then stained with Brilliant Ultra Violet anti-mouse CD69 (BD Biosciences, Franklin Lakes, NJ), PE/Cyanine7 anti-mouse IL-17A (Biolegend), PerCP/Cyanine5.5 anti-mouse IFNγ (Biolegend), eFluor 450 anti-mouse TNFα (eBioscience), PE-CF594 anti-mouse IL-2 (BD Biosciences), and Brilliant Violet 650 anti-mouse IL-4 (BD Biosciences) and analyzed via flow cytometry as we have described in our previous studies (*27, 31, 35*) on the BD FACSymphony A5.

### B cell phenotypes

Splenocytes isolated from above mice were directly subjected to B cell panel staining without rF1V stimulation. Briefly, splenocytes (2×10^6^ cells) were blocked with anti-mouse CD16/32 antibodies (Biolegend) followed by staining with fixable viability dye eFluor 780 (Invitrogen), FITC anti-mouse CD19 (Biolegend), Alexa Fluor 700 anti-mouse CD3 (Biolegend), Brilliant Violet 650 anti-mouse CD138 (BD Biosciences), PE-Cy7 anti-mouse CD38 (Biolegend), PerCP/Cyanine 5.5 anti-mouse GL7 (Biolegend), Brilliant Violet 510 anti-mouse IgD (Biolegend), Brilliant Violet 421 anti-mouse CD80 (Biolegend), Brilliant Violet 605 anti-mouse CD73 (Biolegend), and PE anti-mouse PDL2 (Biolegend) and analyzed via flow cytometry on the BD FACSymphony A5. Flow cytometry data was prepared via FlowJo software v10.10.

### *In vivo* treatment with antibodies

For IFNγ and TNFα depletion, mice were injected intraperitoneally (i.p.) in the lower right quadrant with anti-IFNγ or anti-TNFα monoclonal antibodies (mAb; 500 µg/mouse/treatment; BioX-Cell, NH; Clone R4-6A2 or XT3.11, respectively) once per day for three days, *i.e.* 24h before challenge, the day of challenge, and 24h after challenge. The corresponding isotype control antibodies were similarly used (IgG1; BioXCell; Clone HRPN).

Statistical analysis.

One-way or two-way analysis of variance (ANOVA) with Tukey’s *post hoc* test was used for data analysis. Animal studies were analyzed using Kaplan-Meier with Mantel-Cox tests. All *in vitro* studies were performed in triplicate while two biological replicates of animal studies were performed.

## One Sentence Summary

IFNγ is dispensable in providing protection against pneumonic plague in mice by our patented live-attenuated and adenovirus-based subunit vaccines.

## Acknowledgements

We would like to thank Dr. Vladimir Motin for providing the YscF antigen in the antibody studies; Meredith Weglarz of the UTMB Flow Cytometry Core; and Dr. Robert Abbott for his generous contribution of B cell flow panel recommendations. Methodology figures created with Biorender.com (2024).

## Author contributions

Conceptualization: EKH, JS, AKC

Methodology: EKH, JS, AKC

Investigation: EKH, JS, PBK, BHN, AKC

Visualization: EKH

Funding acquisition: AKC

Formal Analysis: EKH

Supervision: AKC

Writing – original draft: EKH, JS, AKC

Writing – review & editing: AKH, JS, AKC

## Data and materials availability

All data are available in the main text.

